# Two accessory proteins govern MmpL3 mycolic acid transport in mycobacteria

**DOI:** 10.1101/581447

**Authors:** Allison Fay, Nadine Czudnochowski, Jeremy Rock, Jeffrey R. Johnson, Nevan J. Krogan, Oren Rosenberg, Michael S. Glickman

**Author notes:** Correspondence to: Michael S. Glickman MD, Immunology Program, Sloan Kettering Institute, 1275 York Ave, New York, NY 10065, 6468882368.

## Abstract

Mycolic acids are the signature lipid of mycobacteria and constitute an important physical component of the cell wall, a target of mycobacterial specific antibiotics, and a mediator of *M. tuberculosis* pathogenesis. Mycolic acids are synthesized in the cytoplasm and are thought to be transported to the cell wall as a trehalose ester by the MmpL3 transporter, an antibiotic target for *M. tuberculosis*. However, the mechanism by which mycolate synthesis is coupled to transport, and the full MmpL3 transport machinery, is unknown. Here we identify two new components of the MmpL3 transport machinery in mycobacteria. The protein encoded by *MSMEG_0736*/*Rv0383c* is essential for growth of *M. smegmatis* and *M. tuberculosis*, is anchored to the cytoplasmic membrane, physically interacts with and colocalizes with MmpL3 in growing cells, and is required for trehalose monomycolate transport to the cell wall. In light of these findings we propose Msmeg_0736/Rv0383c be named “TMM transport factor A”, TtfA. The protein encoded by *MSMEG_5308* also interacts with the MmpL3 complex, but is nonessential for growth or TMM transport. However, MSMEG_5308 accumulates with inhibition of MmpL3 mediated TMM transport and stabilizes the MmpL3/TtfA complex, indicating that it stabilizes the transport system during stress. These studies identify two new components of the mycobacterial mycolate transport machinery, an emerging antibiotic target in *M. tuberculosis*.

## Introduction

Mycobacteria have a complex cell wall, which is crucial for maintaining cell integrity, protects against environmental stress, provides a barrier against access of potentially harmful molecules into the cell, and plays a critical role in mycobacterial pathogenesis. The cell wall of mycobacteria is comprised of the common bacterial cell wall glycopolymer, peptidoglycan, external to the cytoplasmic membrane, as well as an additional covalently attached glycopolymer layer comprised of arabinogalactan. Arabinogalactan bridges peptidoglycan and mycolic acids, the signature long chain lipid of mycobacteria. Arabinogalactan esterified mycolates constitute the inner leaflet of the outer membrane bilayer, with the outer leaflet being comprised of hydrophobically associated complex lipids including trehalose dimycolate, sulfolipids, lipomannan and lipoarabinomannan. This outer membrane increases both the complexity of the cell wall structure and its hydrophobicity. The enzymatic steps of the arabinogalactan and mycolate precursor biosynthesis have been well described and are the targets of several antimycobacterials, including isoniazid and ethambutol [1–3].

Mycolic acid synthesis begins with FASI system that produces C16-C18 and C24-C26 fatty acids. The FASII system then extends these products to produce the long meromycolate chains that are the substrates for the polyketide synthetase, Pks13. Pks13 catalyzes the final condensation step to produce α-alkyl β-ketoacids (C60-C90) which are then acetylated and transferred to the 6 position of trehalose [4,5]. Mono-α-alkyl β-ketoacyl trehalose is then reduced by CmrA to trehalose monomycolate (TMM) presumably in the inner leaflet of the cytoplasmic membrane [6,7]. TMM can then be modified by non-essential mycolic acid methyltransferases to produce cyclopropane rings and methyl branches, and in the case of *M. tuberculosis* these modifications alter host-mycolic acid responses [8–14].

After synthesis, TMM must be transported across the cytoplasmic membrane to reach the cell wall; this step is known to require the MmpL3 transporter [7,15–17]. The MmpLs [**<u>M</u>**ycobacterial **<u>m</u>**embrane **<u>p</u>**rotein, **<u>L</u>**arge) are multi-substrate transporters of the Resistance-Nodulation-Division (RND) class that usually act as homotrimers and are exporters of molecules from the outer leaflet of the plasma membrane, to, or through the outer membrane. In *Mtb* they include lipid and fatty acid transporters of virulence-associated lipids across the cell envelope. Transport is driven by downhill movement of H^+^ in response to the electrochemical H^+^ gradient (Δ*μ̃*_H_+) across the plasma membrane. MmpL3 is the only MmpL protein that is essential *in vitro*, though mutations in several other MmpLs severely compromise virulence in infection models [7,15–18]. Mutational analyses and ‘transposon-site-hybridization’ (TraSH) revealed MmpL3 is essential for *Mtb* viability *in vitro* [19] and *in vivo* [20], and several inhibitors of MmpL3 are already in clinical development, among them are a set of diamine-indole-carboxamides [21–23] including Novartis NITD-304, and the pyrrole BM212 [24].

Genetic, pharmacologic, and biochemical studies strongly indicate that the MmpL3 transporter is the TMM flippase. MmpL3 has been shown to have flippase activity in spheroplast assays [25] and genetic depletion leads to growth arrest and loss of TMM transport [26,27]. Recent crystal structures of MmpL3 suggest potential mechanisms of TMM transport [28]. However, the full mechanisms linking TMM biosynthesis to MmpL3 transport, and the full set of cofactors used by MmpL3 to transport TMM, are unknown.

MmpLs share close homology with other bacterial RND proteins, typified by the acridine resistance complex (AcrB) transporter that is involved in the efflux of hydrophobic small molecules from or through the periplasm of *E. coli*. Like AcrB, the MmpLs are thought to be localized to the inner membrane [29]. However AcrB does not act alone: it additionally forms the core of a comprehensive secretion system that traverses both the inner and outer membrane of the cell envelope in Gram negative bacteria, allowing the AcrB substrates to bypass the periplasm [30]. To form this membrane spanning system, AcrB interacts with a periplasmic coupling protein called AcrA (or more generally, the Membrane Fusion Protein (MPF)), which in turn links to an outer membrane channel called TolC (or more generally, the Outer Membrane Protein (OMP)) [31–33] The mechanism of bacterial RND transporters is thought to be highly conserved and involves the engagement of the Proton Motive Force (Δ*μ̃*_H_+) to drive drugs, ions and other small molecules from the periplasm across the outer membrane through the MPF, thus preventing the entrance of toxic substances into the bacterial cytoplasm[34–36]. We thus have hypothesized that MmpL3 acts in concert with other mycobacterial proteins, but no MmpL3 associated proteins have been identified. Here we describe two previously unknown cofactors for MmpL3, one of which is required for TMM transport, and one of which is stress inducible and stabilizes the MmpL3 complex.

## Results

### MmpL3 is stably associated with two proteins of unknown function, MSMEG_0736 and MSMEG_5308

In order to discover stable binding interactions with MsMmpL3 *in situ*, we devised a native, stringent, affinity purification. MsMmpL3 was fused to a flexible linker connecting the C-terminus of MmpL3 to monomeric superfolder GFP (msfGFP) at the native chromosomal locus of MmpL3. As *mmpL3* is an essential gene, the normal growth rate of this strain suggests that fusion did not disrupt the essential function of the protein. Cell membranes were collected and solubilized with the mild detergent n-Dodecyl β-D-maltoside (DDM). Anti-GFP nanobodies covalently linked to a magnetic bead were incubated with detergent-solubilized membranes and then extensively washed with 0.2% DDM containing buffer. Co-purifying proteins were identified via shotgun mass spectrometry (Fig 1). One of the most abundantly co-purifying proteins was a protein of unknown function, MSMEG_0736 (Table 1 and Table S1A,B). In contrast, pulldown of MmpL10, another MmpL transporter, did not copurify MSmeg_0736 or any proteins in common with MmpL3 (Table S1A,B). To validate this interaction, we created a strain in which a msfGFP was fused to MSMEG_0736. When MSMEG_0736-msfGFP was purified from detergent solubilized membranes under the same conditions, the most abundantly copurified protein was MsMmpL3 (Table 1 and Table S1A,B). In a control experiment using MSMEG_0410 (MmpL10) fused to msfGFP as a bait, neither MSMEG_0736, MSMEG_0250 or MSMEG_5308 were co-purified (Table 1 and Table S1A,B). In a biological replicate of the MSmeg_0736 pulldown, we confirmed the identity of the prominent band at approximately 100 kDa as MmpL3 (Fig 1B, Table S2). As MSMEG_0736 interacts with MmpL3, and evidence we will present in this paper shows MSMEG_0736 is required for TMM transport, we propose MSMEG_0736 be named “TMM transport factor A”, TtfA.

**Figure 1.**
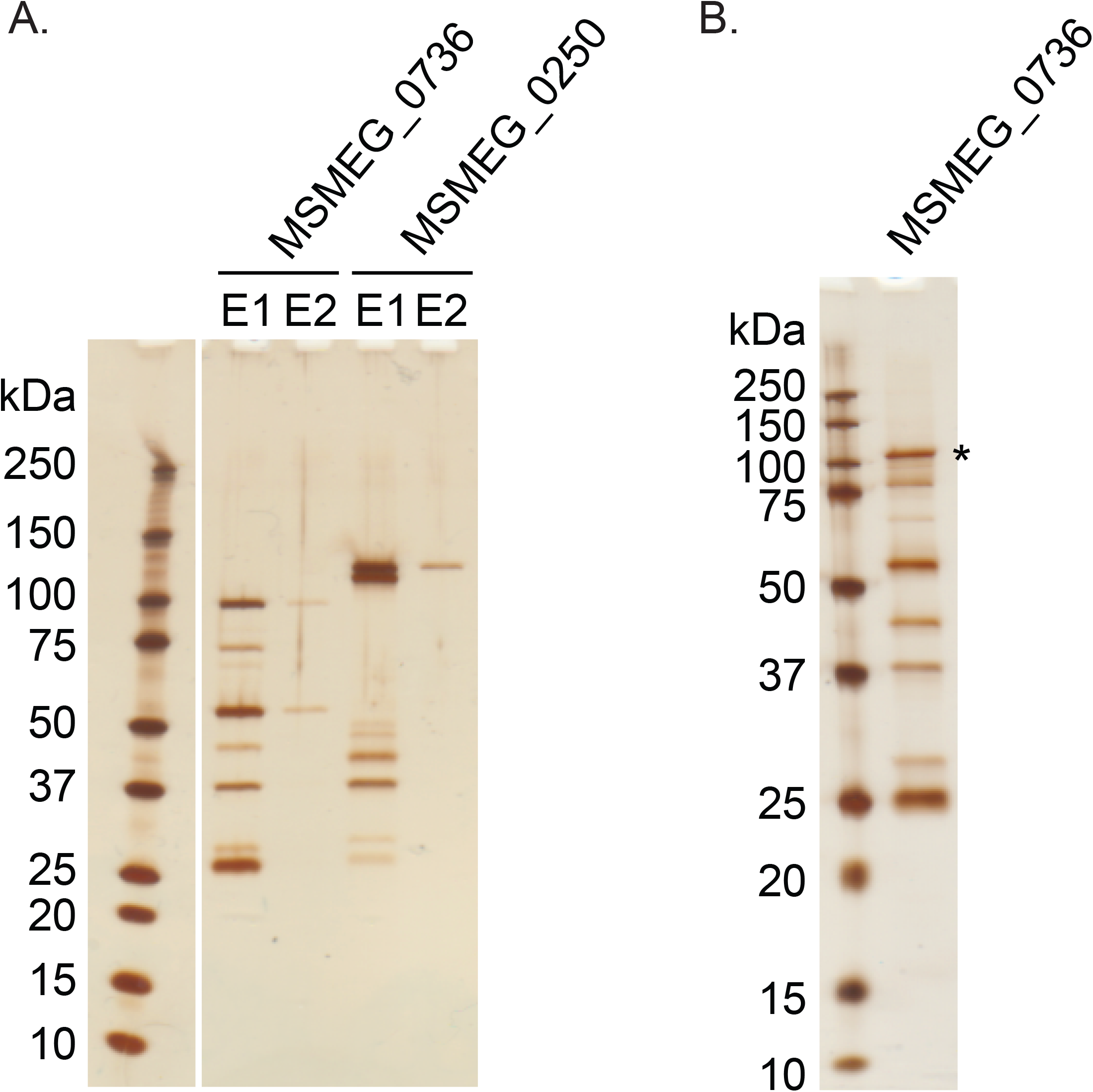
MmpL3 and MSMEG_0736 form a complex. **A.** Silver stained SDS PAGE gel of elutions from GFP-Trap columns loaded with lysates from *M. smegmatis* expressing MSMEG_0736-msfGFP or MmpL3-msfGFP (MSmeg_0250). See Table 1 for protein identifications. **B.** Silver stained SDS PAGE gel of the first elution from a GFP-Trap column loaded with a lysate from *M. smegmatis* expressing MSMEG_0736-msfGFP. The band corresponding to the molecular weight of MmpL3, indicated with an asterisk, was excised and subjected for mass spectrometry analysis and identified as MmpL3 (see methods and Table S2).

**Table 1.** MSMEG_0736 and MSMEG_0250 protein-protein interactions. Shown is the number of unique peptides detected in the GFP-Trap eluates using MSMEG_0736-msfGFP or MSMEG_0250-msfGFP as bait. Only the three proteins with the highest peptides are shown.

| Bait | ProtID | Name | Num unique peptides |
| --- | --- | --- | --- |
| MSMEG_0736 | A0QP27 | MSMEG_0250 | 35 |
|  | A0QQF4 | MSMEG_0736 | 22 |
|  | A0R316 | MSMEG_5308 | 14 |
| MSMEG_0250 (MmpL3) | A0QP27 | MSMEG_0250 | 57 |
|  | A0QQF4 | MSMEG_0736 | 11 |
|  | A0R316 | MSMEG_5308 | 15 |

Analysis of MsTtfA and MsMmpL3 copurifying proteins identified by anti-GFP nanobody purification showed a third complex member found in both pulldowns, the protein encoded by *MSMEG_5308*. This seven bladed beta-propeller protein has a homolog in *M. tuberculosis*, Rv1057, that has been shown to be non-essential, although Mtb lacking Rv1057 fails to properly secrete ESAT-6 and replicated poorly in macrophages [37]. The Rv1057 gene has been shown to be under control of two two-component systems involved in sensing cell stress, MprAB and TcrRS, as well as the envelope stress responsive sigma factor SigE [38–40]. Rv1057 was also reported to be the most transcriptionally induced gene in response to MmpL3 depletion [41], suggesting a connection to MmpL3 function.

### TtfA is essential for growth of M. smegmatis and M. tuberculosis in vitro

The *M. tuberculosis* H37Rv homolog of TtfA is Rv0383c. *rv0383c* was predicted to be an essential gene in H37Rv based on transposon mutagenesis [19,42], but its essentiality in *M. smegmatis* and *M. tuberculosis* is unknown and its molecular function obscure. With no predicted protein domains or homologs of known function, confirmation of its essentiality in both organisms was the first step to analyze its function. To test the essentiality of *ttfA* in *M. smegmatis*, we generated a merodiploid strain in which a second copy of *ttfA* was integrated in the chromosome. We then deleted the endogenous coding sequence, so that the only a single copy of *ttfA* remained at the *attB* site. We then attempted to remove the second copy of *ttfA* from *attB* by marker exchange with either a vector or a plasmid encoding TtfA and conferring kanamycin resistance, pAJF792 [43]. Only transformation with the plasmid encoding TtfA yielded transformants that were kanamycin resistant and streptomycin sensitive. Similar results were obtained with a plasmid encoding TtfA from *M. tuberculosis* (Fig 2A). This inability to remove *ttfA* from *attB* in our Δ*ttfA* strain suggested that *ttfA* was required for growth of *M. smegmatis* (Fig 2A). To further assess the essential role of MsTtfA, we generated CRISPR interference (CRISPRi) strain that allows anhydrotetracycline (ATc) inducible knockdown [44]. Growth inhibition by gene knockdown was visualized by spotting 10-fold serial dilutions on plates with and without ATc, MsTtfA depletion led to an ATc dependent growth defect not seen in the non-targeting control (Fig 2A). Gene knockdown of *ttfA* in *M. smegmatis* also led to cessation of growth in liquid media between 9 and 12 hours post induction with ATc (Fig 2B). To test whether TtfA was essential in *M. tuberculosis*, we attempted to knockout the gene using a temperature sensitive phage and were unsuccessful, suggesting essentiality. We then generated three *ttfA* targeting CRISPRi strains with independent guide RNAs. Gene knockdown of *ttfA* in *M. tuberculosis* with all three guide RNAs all led to cessation of growth in liquid media after three days after induction with ATc, indicating that TtfA is essential for *M. tuberculosis* growth *in vitro* (Fig 2C).

**Figure 2.**
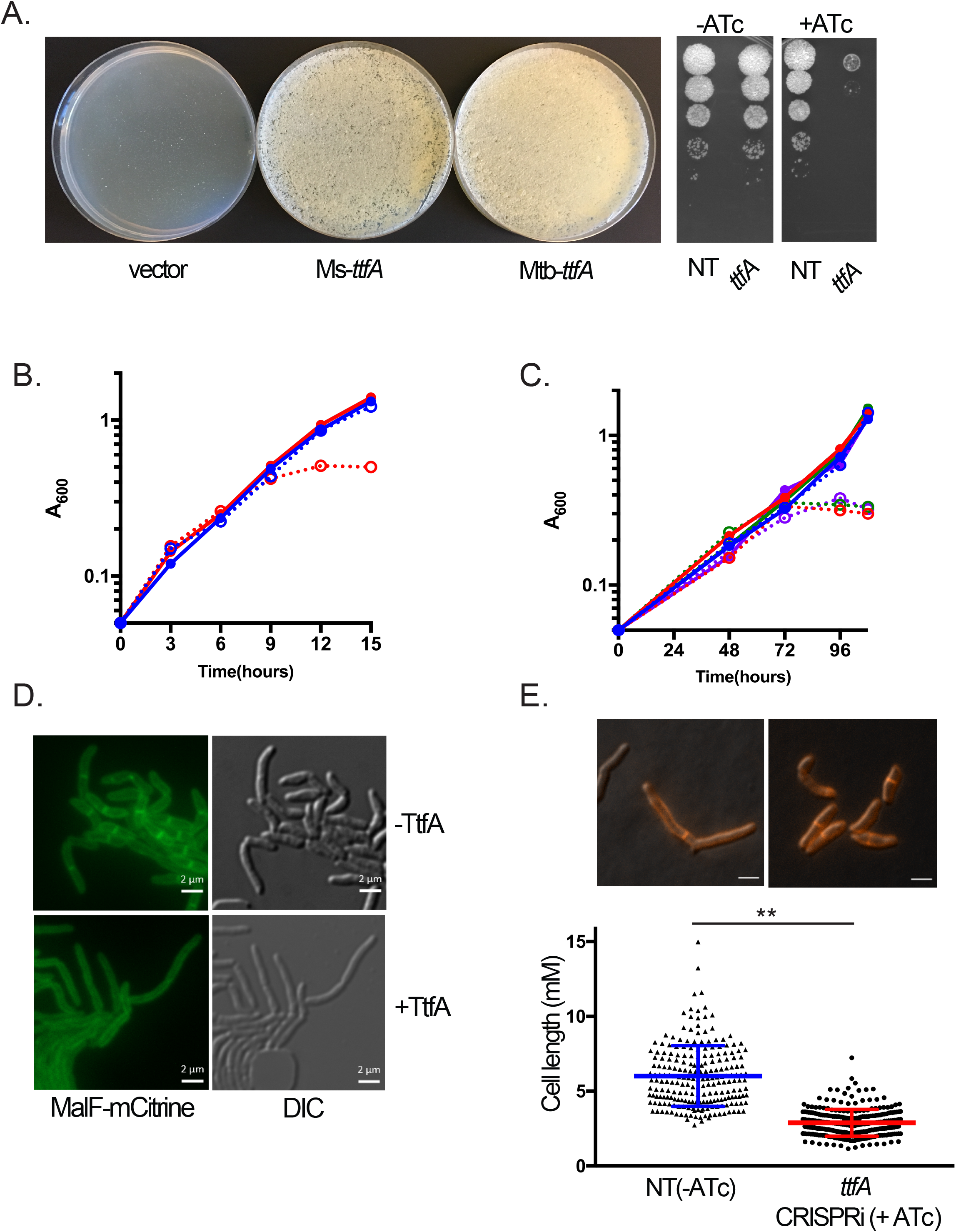
MsTtfA /Mtb TtfA are required for mycobacterial growth and cell elongation. (A) (Left) *M. smegmatis* strains carrying deletion in chromosomal *ttfA* and a copy of *ttfA* at the *attB* phage integration site were subjected to marker exchange with *attB* integrating vectors. Δ*ttfa attB*::*ttfA strep* (MGM6414) transformed with pMV306kan (vector), pAJF792 (encoding MsTtfA) or pAJF793 (Mtb TtfA) are shown on Kanamycin agar. (Right) 10-fold dilutions of *M. smegmatis* carrying ATc inducible CRISPRi non-targeting control (NT, MGM6418) or *ttfA* (MGM6419) on agar media with and without ATc. (B) Growth curve of non-targeting (MGM6418, blue) and *ttfA* targeting (MGM6419, red) CRISPRi *M. smegmatis* strains grown in uninduced (solid, closed circles) and ATc induced (dashed, empty circles) conditions. (C) Growth curve of non-targeting (MGM6715, blue) or three distinct Mtb *ttfA* targeting CRISPRi *M. tuberculosis* strains (MGM6675, red; MGM6677, green; MGM6679, purple) grown in uninduced (solid, closed circles) and ATc induced (dashed, empty circles) conditions. (D) Fluorescence microscopy of a Ms *ttfA* targeting CRISPRi strain marked with MalF(1,2)-mCitrine (MGM6433) 15 hours post CRISPRi induction with ATc (top, -TtfA) or, uninduced control at 15 hours (+TtfA, bottom). YFP (left) and DIC (right) image shown. White bar indicated in bottom of panel image is 2 micron. Exposure times for YFP 250ms, 40% LED. (E) Loss of TtfA leads to short cells. Cell lengths of non-targeting (MGM6418, graph:blue, triangles) and TtfA targeting (MGM6419, red, squares) CRISPRi strains induced for 12 hours. Representative DIC /FM 4-64 images used for quantitation shown above the graph.

To examine the morphologic changes that accompany growth arrest during loss of MsTtfA, we depleted the protein using CRISPRi and tracked morphological changes using a MalF(1,2)-mCitrine expression strain that uniformly labels the cell membrane. Time-lapse microscopy indicated that growth arrest without MsTtfA was characterized by shorter, misshapen cells (Fig 2D, Movies S1, S2). Quantitation of cell length revealed that MsTtfA depleted cells were significantly shorter (2.88± 0.89 *μ*m) as compared to control cells (6.00±2.03 *μ*m) (Fig 2E). The short cell phenotype suggested that MsTtfA might be required for cell elongation. These data indicate that TtfA is essential for mycobacterial viability and that the function of this gene is conserved between fast and slow growing mycobacteria.

### MSMEG_0736 localizes to poles and septa

The predicted protein encoded by MsTtfA contains a predicted N-terminal transmembrane domain from amino acids 2-24, indicating that it is either a transmembrane or secreted protein. To determine the localization and topology of MsTtfA, we assessed the *in vivo* functionality of mCitrine fused at the N or C-terminus. Marker exchange with a plasmid encoding MsTtfA-mCitrine yielded kanamycin resistant, streptomycin sensitive transformants in similar numbers to pAJF792, encoding the wildtype gene, indicating the C-terminal fusion is functional. In contrast, the plasmid encoding an N-terminal mCitrine fusion did not yield kanamycin resistant, streptomycin sensitive transformants, indicating that this fusion failed to complement for essential function.

We next localized MsTtfA using live cell fluorescence microscopy. The C-terminal mCitrine fusion protein produced fluorescent signal at the cell poles and septa (Fig 3A, and Movie S3). It has been previously reported that mCitrine does not fluoresce when localized in the periplasm, suggesting that the C-terminal domain of MsTtfA is localized in the cytoplasmic side of the membrane [45]. We then generated an MsTtfA C-terminal fusion to msfGFP by recombination such that the fused copy was expressed from its endogenous locus and was the only copy, guaranteeing functionality. The resulting MsTtfA-msfGFP strain demonstrated fluorescent signal exclusively at cell poles and septa (Movie S4). Fractionation of cell free supernatants showed no detectable MsTtfA-msfGFP in the supernatant (Fig 3B), suggesting that the protein is not secreted. Fractionation of the cell lysate showed that MsTtfA-msfGFP localized in the Trition-X100 soluble fraction, similar to a membrane protein control FtsY, but not the soluble fraction marked by cytosolic RNAPβ, supporting that MsTtfA is membrane anchored, is not secreted, and has a cytoplasmic C-terminus.

**Figure 3.**
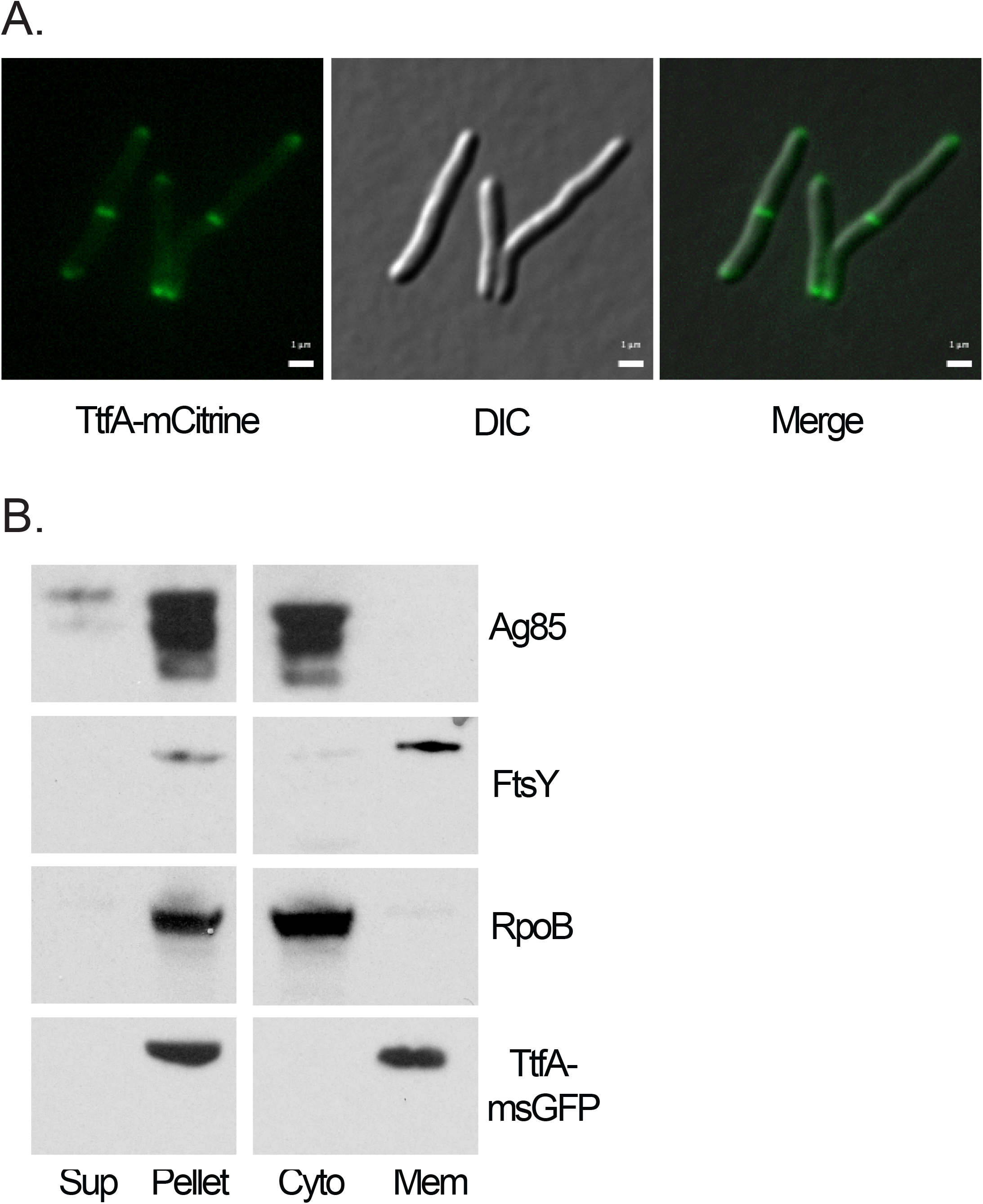
TtfA is a membrane protein that localizes to poles and septa. (A) *M. smegmatis* TtfA-mCitrine expression strain (MGM6423) imaged during logarithmic growth. YFP (left), DIC (middle), and Overlay (right) image shown. White bar is 1 micron indicated in bottom of panel image. Exposure times for YFP 1s, 75% LED. (B) Localization of TtfA-msfGFP by cellular fractionation. Cell free supernatant and cell pellet fractions (left) and cytoplasmic and membrane fractions (right) probed for secreted protein Ag85 (top), membrane protein FtsY (top, middle), cytoplasmic protein RpoB (bottom, middle), and GFP for TtfA-msfGFP (bottom).

### The essential portion of TtfA is conserved among mycolate producers

To further delineate the functional domains of the protein, we examined the conservation of the protein sequence across homologs. BLAST searches identified homologous predicted proteins among mycolate producing organisms (Fig S1). Alignments of these homologs suggested that amino acids 1 through approximately 205 were well conserved, with poor conservation in the C-terminal 73 amino acids (Fig S1). The C-terminal 73 amino acids are also predicted to be disordered [46]. This lack of conservation at the C-terminus was also apparent in the alignment with the MtbTtfA, which we demonstrate above is functional in *M. smegmatis* (Fig 2A). To assess the functional contribution of these conserved regions, we generated MsTtfA truncations fused at the C-terminus to msfGFP and assessed the ability of these truncations to complement the essential function by marker exchange. Only the plasmid encoding amino acids 1-205 yielded kanamycin resistant, streptomycin sensitive transformants, indicating that amino acids 1-205 were essential (Fig S2A).

After confirming that all of these truncations accumulate as stable proteins at their predicted sizes when expressed in wild type *M. smegmatis* (Fig S2A), we localized each truncation by fluorescence microscopy. MsTtfA(1-205aa)-msfGFP localized to poles and septa in a pattern similar to the full-length protein (Fig S2B), indicating that the poorly conserved C-terminus is not required for essential function or proper localization. However, truncations shorter than 205AA, which did not complement essential function, also failed to localize to poles and septa, indicating that the first 205AA of the protein, including the N terminal transmembrane domain, are required for proper localization and that this localization is tightly linked to its essential function.

### The N-terminus of TtfA is required for interaction with MmpL3

To determine the regions of MsTtfA required for interaction with MmpL3, we immunopurified MsTtfA truncations fused to msfGFP when coexpressed with MmpL3-mCherry. MsTtfA-msfGFP was purified from DDM detergent solubilized lysates with GFPTrap resin. Unfused msfGFP did not coprecipitate MmpL3-mCherry, whereas full-length MsTtfA-msfGFP copurified with MmpL3-mCherry (Fig 4A). All truncations were visible at comparable levels in DDM solubilized lysates at their predicted sizes (Fig 4B). However, only MsTtfA(1-205aa)-msfGFP copurified with MmpL3-mCherry, quantitatively similar to full-length MsTtfA-msfGFP (Fig 4B). However, loss of any segment of MsTtfA within the first 205AA abolished interaction with MmpL3. These results demonstrate an exact correlation between the ability of MsTtfA to interact with MmpL3 and the essential function of this protein, suggesting that the essentiality of TtfA may be due to a role as an MmpL3 cofactor.

**Figure 4.**
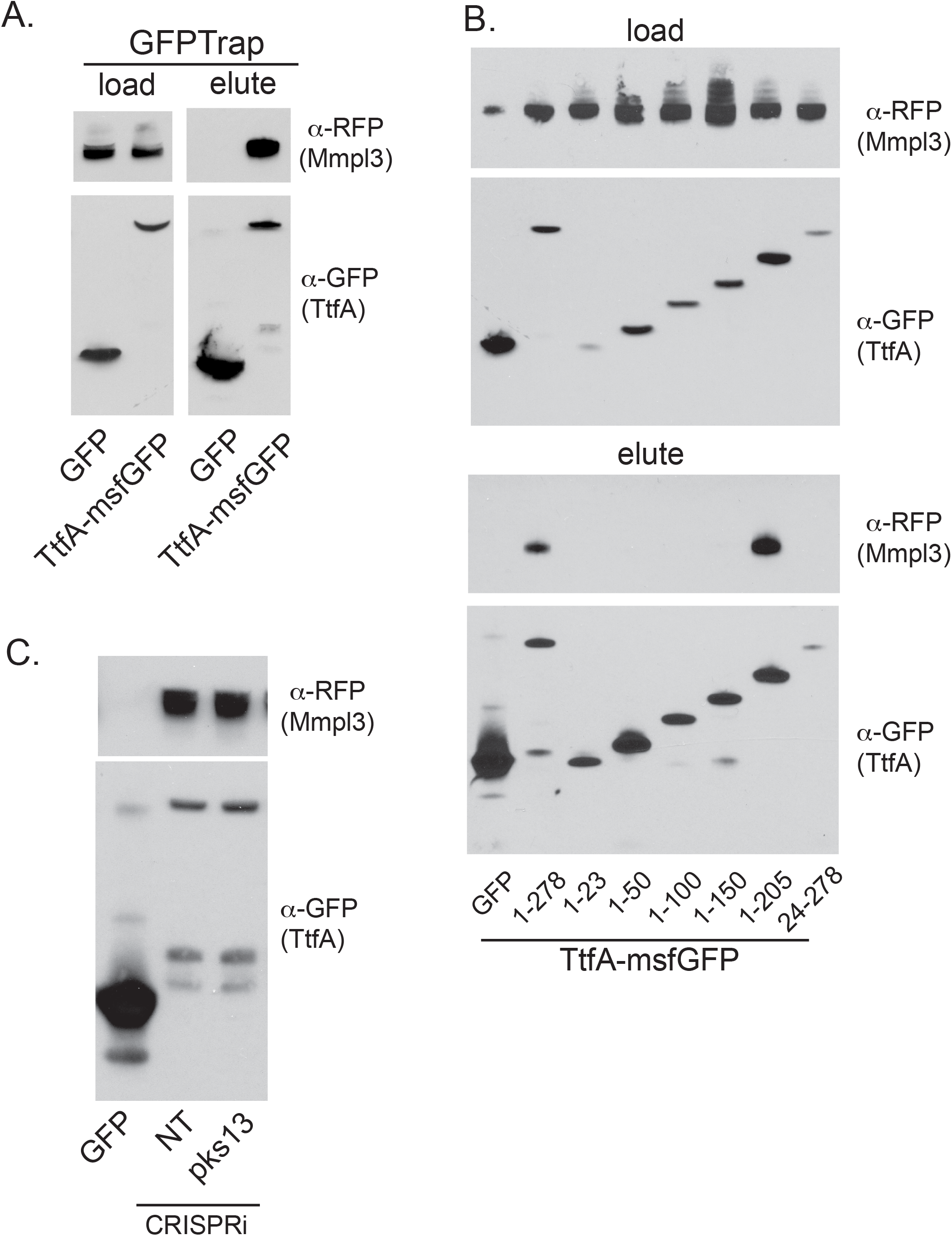
TtfA and MmpL3 form a complex *in vivo* via the essential region of TtfA and independently of TMM synthesis. (A) DDM solubilized *M. smegmatis* lysates (left) and GFPTrap eluates (right) of msfGFP expressing control (MGM6828) and TtfA-msfGFP (MGM6815) both co-expressing MmpL3-mCherry and probed with anti-RFP (top) and anti-GFP (bottom). (B) The essential region of MsTtfA is necessary and sufficient for MmpL3 interaction. DDM solubilized lysates (top) and GFPTrap eluates (bottom) of msfGFP control (MGM6828), full-length TtfA-msfGFP (1-278, MGM6829), or TtfA-msfGFP truncations (1-23:MGM6826, 1-50:MGM6823, 1-100:MGM6827, 1-150:MGM6824, 1-205:MGM6822, 24-278:MGM6825) co-expressing MmpL3-mCherry and probed with anti-RFP (top) and anti-GFP (bottom). (C) The MsTtfA-MmpL3 interaction is independent of mycolate synthesis. GFPTrap eluates of MmpL3-mCherry expression strains co-expressing msfGFP control (MGM6828) or TtfA-msfGFP with either control CRISPRi for (NT, MGM6816) or *pks13* (MGM6817) depleted for 6 hours with ATc. Top panel is probed for MmpL3-mCherry with anti-RFP and bottom with anti-GFP.

### The MmpL3 and MSMEG_0736 complex and its localization is independent of TMM biosynthesis

To examine whether the TtfA-MmpL3 interaction requires TMM synthesis, the substrate of the MmpL3 flippase, we depleted Pks13, the TMM synthetase in *M. smegmatis* [47,48]. Depletion of Pks13 in the MmpL3-mCherry/TtfA-msfGFP caused growth arrest after 6 hours of induction, indicative of depleting essential Pks13 (Fig S3) However, Pks13 depletion did not affect levels of either TtfA-msfGFP or MmpL3-mCherry in DDM solubilized lysates (data not shown), nor did depletion of Pks13 have any effect on the TtfA-msfGFP-MmpL3-mCherry complex (Fig 4C). These results indicate that active TMM biosynthesis is not required for TtfA-MmpL3 complex formation.

MmpL3-GFP has been previously reported to localize to cell poles and septa [49], a finding we confirm with our MmpL3-msfGFP strain, which localizes the MmpL3 protein to poles and septa (Movie S5). This pattern is very similar to the pattern observed with TtfA-msfGFP (Movie S4). To colocalize MmpL3 and TtfA we again utilized strains co-expressing mCherry and msfGFP fusions to TtfA and MmpL3. By fluorescence microscopy, MmpL3 and TtfA strongly co-localized to cell poles and septa (Fig 5A) and were indistinguishable in their localization patterns. Depletion of Pks13 via CRISPRi led to cessation of growth between 6 and 9 hours, but did not affect localization of TtfA-msfGFP or MmpL3-msfGFP, again indicating that TMM synthesis was not required for localization of either protein to the poles or septa (Fig 5B). Taken together, these results strongly indicate that MmpL3 and TtfA form a complex *in vivo* at the site of cell growth.

**Figure 5.**
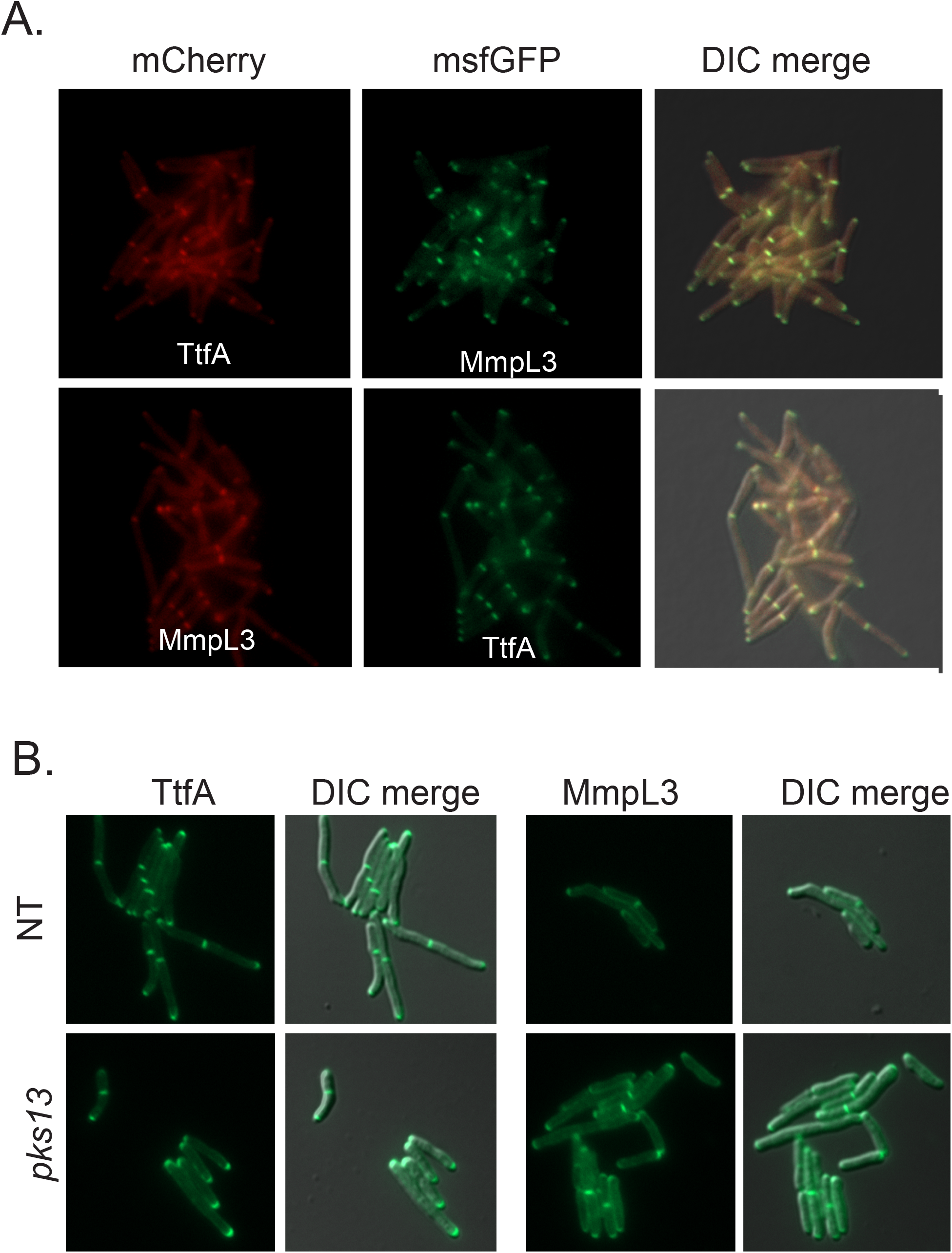
TtfA and MmpL3 co-localize at cell poles and septa independently of TMM synthesis. (A) Localization of MsTtfA-mCherry/MmpL3-msfGFP (MGM6433, top) and MmpL3-mCherry/TtfA-msfGFP (MGM6434, bottom). (B) Localization of TtfA-msfGFP or MmpL3-msfGFP in Pks13 depleted or mock depleted cells.

### TtfA is required for MmpL3 TMM transport in *M. smegmatis* and *M. tuberculosis*

We next assessed whether TtfA is functionally required for TMM flipping. Loss of MmpL3 function via genetic or pharmacologic inhibition results in TMM accumulation and TDM depletion due to the inability of TMM to flip across the cytoplasmic membrane where Antigen85 enzymes process TMM to TDM and arabinogalactan attached mycolic acids [25–27]. To assess the functional role of TtfA in TMM flipping, we utilized CRISPRi strains that depleted TtfA or MmpL3, and a non-targeting control in *M. smegmatis*. 6 hours after knockdown of gene expression, we labeled mycolic acids with ^14^C-acetate and assessed TMM/TDM levels in cell wall organic extracts. Depletion of MmpL3 had the reported effect of ^14^C-TMM accumulation and ^14^C-TDM depletion (Fig 6A), attributable to MmpL3 dysfunction. Depletion of TtfA has a quantitatively similar effect on TMM transport as depletion of MmpL3, as shown by the TDM/TMM ratio in depleted cultures as compared to replete cultures (Fig 6A,B). As a control for essential protein depletion, we depleted the essential DnaK chaperone [43] and found no effect on ^14^C-TMM/^14^C-TDM, indicating that cell arrest by depletion of any essential protein does not alter TMM and TDM levels (Fig S4).

**Figure 6.**
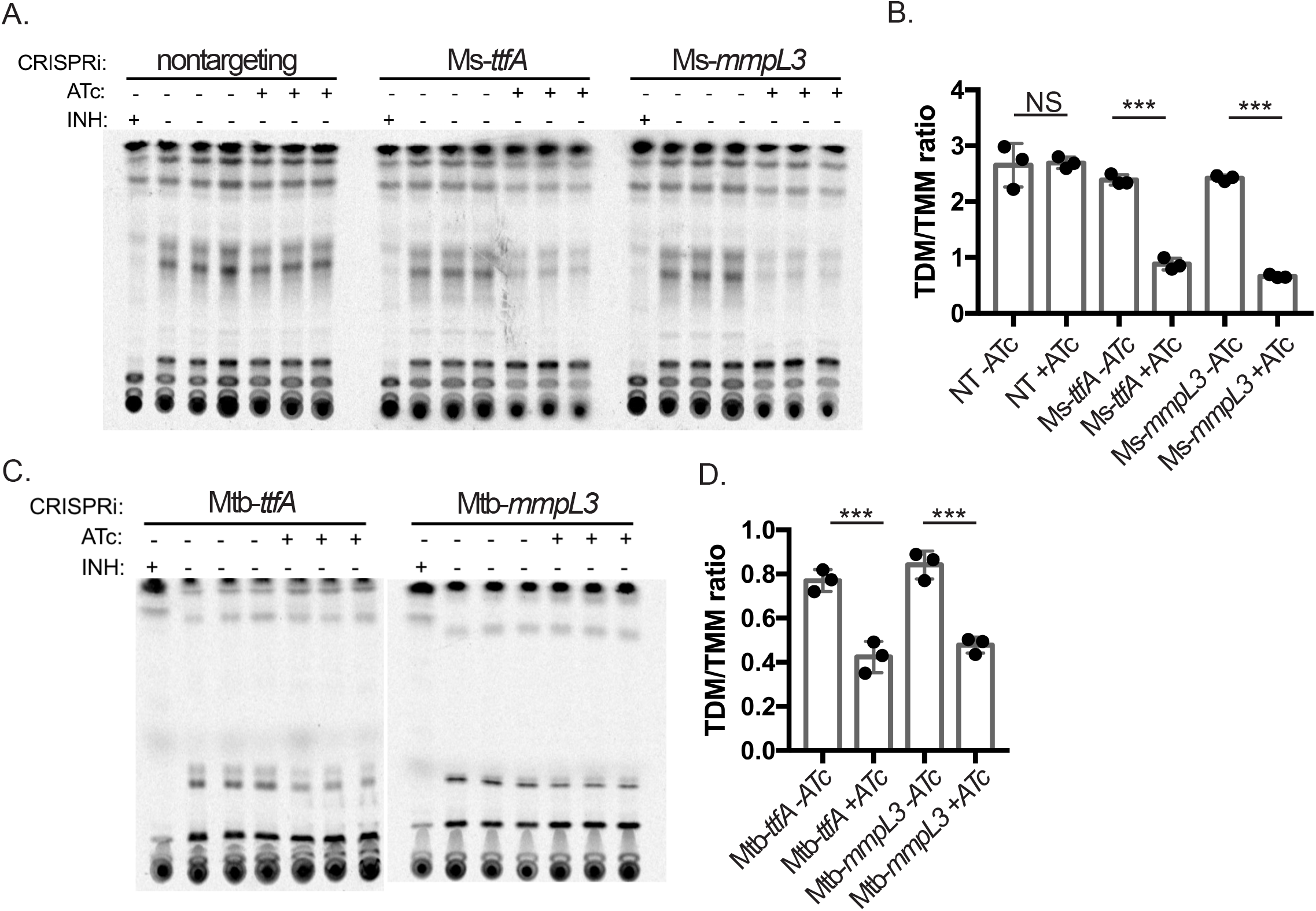
MsTtfA and MtbTtfA are required for TMM transport. (A) TLCs of extractable mycolic acids from three replicate ^14^C-acetic acid labeled *M. smegmatis* cultures carrying CRISPRi targeting guide RNAs (non-targeting, MGM6418), *ttfA* (middle, MGM6419), or *mmpL3* (right, MGM6637) (B) Graph of TDM/TMM ratio for quantitation of TMM and TDM of TLCs in panel A. (C) TLCs of extractable mycolic acids from three replicate ^14^C-acetic acid labeled *M. tuberculosis* cultures depleted for TtfA (left, MGM6675), or MmpL3 (right, MGM6676) (D) Quantitation of TDM/TMM ratio from quantitation of TMM and TDM of TLCs in panel C.***=p<0.01.

We saw similar results in *M. tuberculosis* depleted of TtfA or MmpL3. Either TtfA or MmpL3 depletion impaired TMM transport, with the resulting accumulation of TMM and loss of TDM (Fig 6C,D). These results indicate that loss of the MmpL3 interacting protein TtfA impairs MmpL3 dependent TMM transport in both *M. smegmatis* and *M. tuberculosis*, strongly indicating that TtfA is an essential cofactor in MmpL3 function.

### An additional complex member is responsive to MmpL3 and TtfA depletion and inhibition of flippase activity

MSMEG_5308 was also found to co-purify with both MsTtfA and MmpL3 (Fig 1 and Table 1). To further investigate this MmpL3 complex member, we generated a C-terminal msfGFP fusion to MSMEG_5308 at the chromosomal locus. We then depleted either MmpL3 or TtfA in the MSMEG_5308-msfGFP strain. Either MmpL3 or TtfA depletion, but not non-targeting control, led to accumulation of MSMEG_5308 protein (Fig 7A). In contrast, CRISPRi depletion of Pks13 led to cessation of cell growth after 6 hours of induction, but did not induce MSMEG_5308 accumulation (Fig 7A).

**Figure 7.**
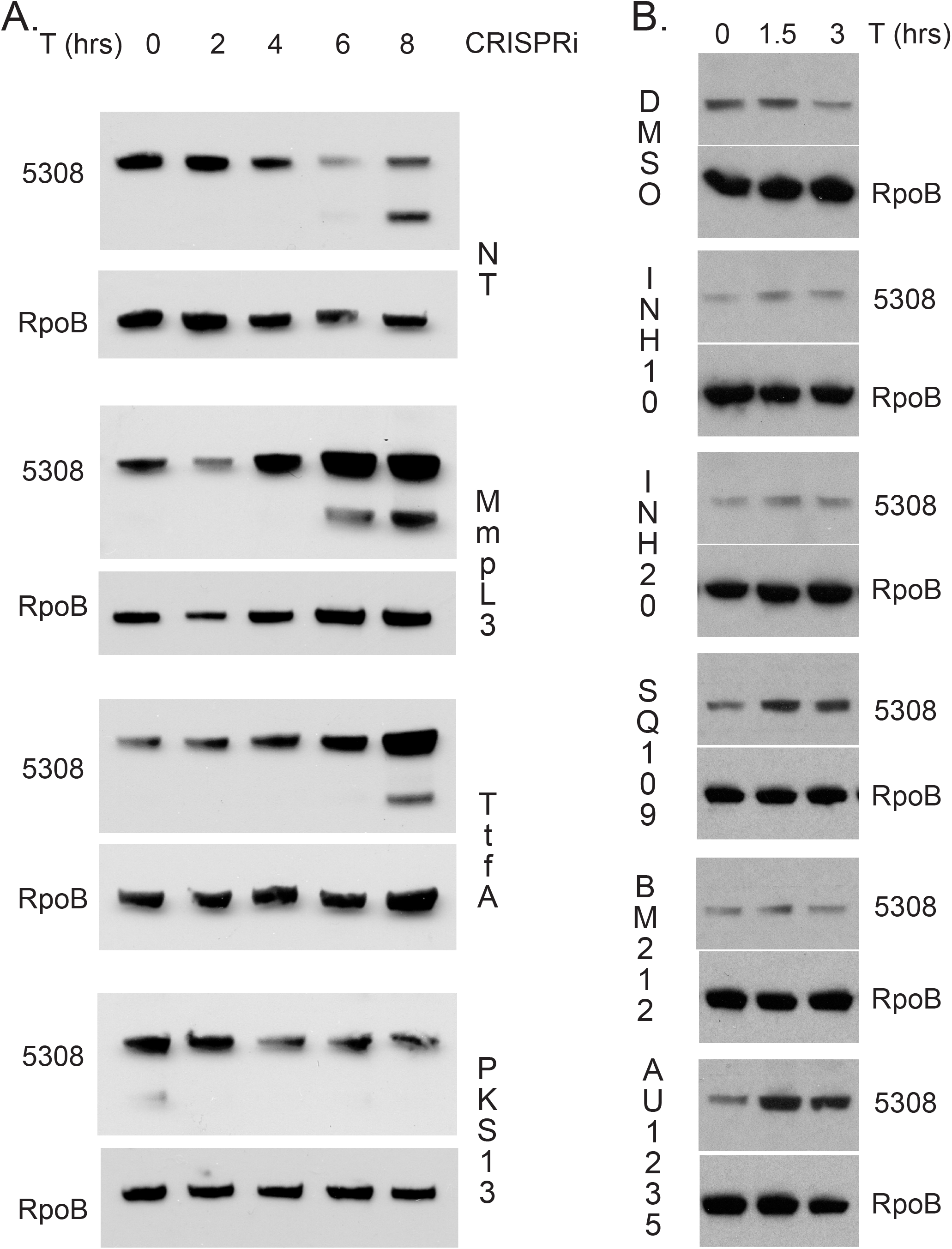
MSMEG_5308-msfGFP accumulates in response to MmpL3 dysfunction. (A) Lysates of MSMEG_5308-msfGFP expression strains with CRISPRi constructs non-targeting control (NT, MGM6766), MmpL3 (MGM6718), TtfA (MGM6717), or Pks13 (MGM6767) (ATc induction at 0, 2, 4, 6, 8 hours) and probed with anti-GFP or anti-RpoB. (B) Lysates of MSMEG_5308-msfGFP expression strain (MGM6681) treated with DMSO, 10μg/ml INH, 20μg/ml INH, 5μM SQ109, 5μM BM212, 5μM AU1235 for 0, 1.5, or 3 hours and probed with anti-GFP and anti-RpoB (loading control)

We further examined the response of MSMEG_5308 to inhibitors of the TMM/TDM pathway, including early mycolate biosynthesis (isoniazid (INH)), and inhibitors targeting late steps in TMM/TDM transport (SQ109, BM212, and AU1235). Resistance mutations to SQ109, BM212, and AU1235 arise in MmpL3, however only BM212 and AU1235 have been shown to directly inhibit MmpL3 flippase activity and all three inhibitors may have effects outside of MmpL3 flippase activity [24–26,50]. Both SQ109 and AU1235 caused MSMEG_5308-msfGFP accumulation at 1.5 and 3 hours, but INH or BM212 (at 5 and 10 μM) had no effect (Fig 7B and data not shown). The lack of accumulation of MSMEG_5308 with INH treatment or Pks13 depletion suggests that MSMEG_5308 does not accumulate in response to loss of TMM or TDM biosynthesis, but rather inhibition of their transport.

### MSMEG_5308-msfGFP localizes to cell poles and septa and stabilizes MmpL3/TtfA interaction

The identification of MSMEG_5308 as an MmpL3/TtfA interacting protein suggested that MSMEG_5308 may co-localize with the MmpL3 complex. Indeed, MSMEG_5308-msfGFP localized to cell poles and septa in a pattern similar to both TtfA-msfGFP and MmpL3-msfGFP by live cell fluorescence microscopy (Fig 8A). To examine the role of MSMEG_5308, we targeted *MSMEG_5308* using CRISPRi and verified efficient knockdown using a MSMEG_5308-msfGFP strain (Fig S5). Depletion of MSMEG_5308 had no impact growth or cell morphology, confirming MSMEG_5308 was not essential in *M. smegmatis* (data not shown).

**Figure 8.**
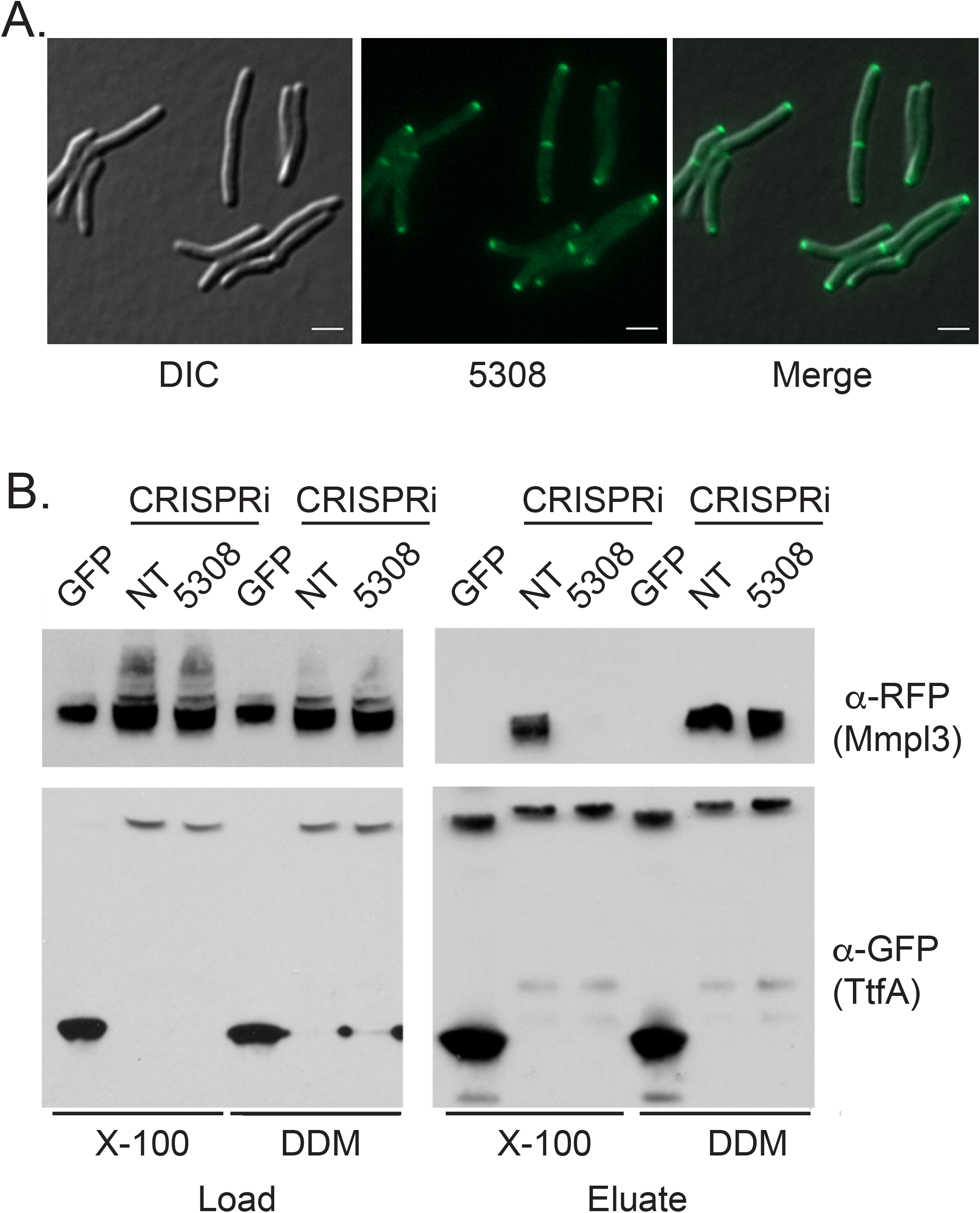
MSMEG_5308 localizes to cell poles and septa and stabilizes the TtfA/MmpL3 interaction. (A) MSMEG_5308-msfGFP expression strain (MGM6681) imaged during logarithmic growth. GFP, DIC and merged images are shown. (B) GFPTrap pulldown of TtfA-msfGFP /MmpL3-mCherry co-expression strains with non-targeting or MSMEG_5308 targeting CRISPRi constructs. Left panel is the input and right panel is the eluate from the GFPTrap column, either in the presence of DDM or Triton X-100.

To assess the effect of MSMEG_5308 on MmpL3/TtfA complexes, we isolated TtfA-msfGFP using anti-GFP nanobodies and probed for MmpL3-mCherry in the presence and absence of MSMEG_5308. In DDM solubilized lysates, TtfA copurified with MmpL3-mCherry in MSMEG_5308 depleted lysates similarly to control cells (Fig 8B). However, in Triton X-100 solubilized lysates, although the MmpL3-TtfA complex was intact when MSMEG_5308 was present, TtfA-msfGFP did not coprecipitate MmpL3-mCherry in the absence of MSMEG_5308 (Fig 8B). These results indicate that MSMEG_5308 is a nonessential member of the MmpL3 complex that is induced by stress and stabilizes the MmpL3-TtfA protein complex.

## Discussion

We have identified two new components of the essential machinery of mycolic acid transport and cell growth in mycobacteria. The MmpL3 transporter was previously known to transport trehalose monomycolate, but its cofactors were unknown. The MmpL3 machinery contains the essential protein TtfA, which we show is required for TMM flipping across the cytoplasmic membrane. A third complex member, MSMEG_5308, while not required for TMM transport, appears to stabilize the MmpL3 complex and is upregulated in response to MmpL3 dysfunction. All three of these proteins localize to cell poles and septa, which are the sites of cell elongation and the previously reported localization sites of early mycolic acid synthetic machinery such as MabA and InhA [49].

TtfA, a protein with no predicted domains of known function, is an essential component of the mycolic acid transport machinery. We have defined the essential portions of TtfA, amino acids 1-205 that includes the N-terminal transmembrane domain but not the poorly conserved disordered C-terminus. Using coprecipitation techniques, we see that truncations that disrupt localization and interaction with MmpL3 fail to support the essential TMM transport function of MmpL3. Our model for the molecular function of TtfA in TMM transport is that the protein links the mycolate biosynthetic machinery to the MmpL3 transporter, possibly by binding directly to TMM. MmpL3 is distinct from several other MmpL proteins in that disruption of the transporter does not inhibit synthesis of the transported lipid. For several MmpL proteins, transport and synthesis are linked. For example, deletion of the sulfolipid transporter MmpL8 abolishes sulfolipid synthesis, rather than simply inhibiting its transport [16,51]. Similarly, MmpL7 is physically and functionally coupled to PDIM biosynthesis [52]. However, the lack of such coupling in the MmpL3 system may suggest that a coupling protein is required to chaperone the transported glycolipid to the transporter, a function we hypothesize for TtfA.

Alternatively, it is possible that that TtfA is a scaffolding protein that nucleates additional essential MmpL3 complex members yet to be elucidated. TtfA has been previously shown to interact with the non-essential vesiclulogenesis regulator VirR in *M. tuberculosis*, that we also find in our purifications of MsTtfA [53].

The second protein we identify in the MmpL3 complex, MSMEG_5308, is a seven bladed propeller protein. This protein structural motif has been previously described to aid in protein-protein interactions though members are functionally diverse [54–56]. In Mtb, the MSMEG_5308 homolog, Rv1057, is responsive to a variety of membrane stresses as well as MmpL3 depletion. Our data indicates that the function of MSMEG_5308 is to stabilize the MmpL3/TtfA complex. We hypothesize that MSMEG_5308 is upregulated during times of membrane stress in order to stabilize MmpL3 complexes and preserve TMM transport and cell wall biosynthesis in conditions that may dissociate the MmpL3 complex.

MmpL3 mediated TMM transport has emerged as an attractive drug target after several high throughput screens identified whole cell active inhibitors that appear to target this transporter. Our identification of two previously unidentified cofactors for MmpL3 will empower future studies to investigate these proteins as drug targets and their potential roles in cellular response and resistance to MmpL3 targeting small molecules. Additionally, future biochemical and structural studies will examine the biochemical and structural organization of this essential mycolic acid transport complex.

## Methods

### Bacterial and DNA manipulations

Standard procedures were used to manipulate recombinant DNA and to transform *E. coli*. *M. smegmatis* strains were derivatives of *mc^2^155* [60]. *M. tuberculosis* strains are derivatives of Erdman. Gene deletions were made by homologous recombination and double negative selection [61]. All strains used in this study are listed in Table S2. Plasmids including relevant features, and primers are listed in Table S3 and S4. *M. smegmatis* and *M. tuberculosis* was transformed by electroporation (2500V, 2.5μF, 1000Ω). All *M. smegmatis* strains were cultured in LB with 0.5% glycerol, 0.5% dextrose (LBsmeg) or 7H9 media. *M. tuberculosis* was growth in 7H9_OADC_. 0.05% Tween_80_ was added to all liquid media. Antibiotic concentrations used for selection of *M. smegmatis* and *M. tuberculosis* strains were as follows: kanamycin 20μg/ml, hygromycin 50μg/ml, streptomycin 20μg/ml. For CRISPRi knockdowns anhydrotetracycline (ATc) was used at 50ng/ml (smegmatis) or 100ng/ml (Mtb).

### Immunoblotting

For protein and epitope tag detection, GFP (Rockland Immunochemicals, Rabbit Anti-GFP polyclonal antibody, 1mg/ml, 1:20,000), mCherry (Rockland Immunochemicals, Rabbit Anti-RFP polyclonal antibody, 1mg/ml, 1:20,000), and RNAP-β (Neoclone, 8RB13 Mouse Anti-E. coli RNAPβ monoclonal, 1:20,000), Ag85 (BEI Resources, Rabbit polyclonal antibody, 1:20000).

### Microscopy

All images were acquired using a Zeiss Axio Observer Z1 microscope equipped with Definite focus, Stage top incubator (Insert P Lab-Tek S1, TempModule S1), Colibri.2 and Illuminator HXP 120 C light sources, a Hamamatsu ORCA-Flash4.0 CMOS Camera and a Plan-Apochromat 100x/1.4 oil DIC objective. Zeiss Zen software was used for acquisition and image export. The following filter sets and light sources were used for imaging: GFP (38 HE, Colibri2.0 470 LED), mCherry (64 HE, Colibri2.0 590 LED). YFP (46 HE, Colibri2.0 505 LED) and FM 4-64 (20, HXP 120 C). For cell staining 100μl of culture was used. A final concentration of 1μg/ml FM 4-64 (Invitrogen) was added. Cells were pelleted by centrifugation at 5000g for 1 minute and resuspended in 50μl of media. For single time point live cell imaging, 7μl of culture was spotted onto a No. 1.5 coverslip and pressed to a slide. For time-lapse microscopy, cells were added to a 1.5% Low melting point agarose LBsmeg pad.For pad preparation, LBsmeg agarose was heated to 65°C and poured into a 17×28mm geneframe (Thermoscientific, AB-0578) adhered to a 25×75mm glass slide. A second slide was pressed down on top and the set-up was allowed to cool at room temperature for 10 minutes. The top slide was removed and the pad was cut and removed so that a 3-4mm strip remained near the center. 2-3μl of *M. smegmatis* culture was added to the pad and a No. 1.5 24×40mm coverglass was sealed to the geneframe. Slides were incubated in stage top incubator at 37°C. For timelaspse microscopy cells were incubated in CellAsic ONIX microfluidic system (plates for bacterial cell culture, B04A) at a flow (psi) of 2.0 and heated at 37°C. Cells were equilibrated in plates at 37°C for 3 hours prior to the start of imaging. Cell length were quantitated using Zeiss Zen software.

### ^14^C-Acetic Acid labeling and TLC

*M. smegmatis* and *M. tuberculosis* cultures were grown and depleted for the following times: M. *smegmatis* CRISPRi 6 hours in ATc-50ng/ml, *M. smegmatis* Tet-DnaK 16 hours without ATc, and *M. tuberculosis* CRISPRi 26 hours in ATc-100ng/ml. For TMM and TDM labeling 1ml of culture was removed and labeled for 1 hour with 1ul for *M. smegmatis* or 16 hours with 2ul for *M. tuberculosis* using [1-^14^C]-Acetic Acid, Sodium Salt (Perkin Elmer, NEC084H001MC, 1 mCi/mL). For INH controls, 20ug/ml INH was added 5 minutes prior to label addition. After incubation cells were harvested by centrifugation at 10,000g for 5 minutes and supernatant was removed. The pellet was resuspended in 500ul chloroform:methanol (2:1) and incubated at 37C for 2 hours. Cells and debris was pelleted at 10,000g for 5 minutes and the supernatant was removed. 10ul of chloroform:methanol extraction was spotted on HPTLC plates and run 3 times in chloroform:methanol:water (90:10:1), then allowed to air dry and imaged using a Phosphor storage cassette and Typhoon Trio (pixel size 200 microns at best sensitivity). ImageJ64 was used to quantitate the radioactive signal.

### Protein expression and purification

Endogenous MSMEG_0736 and MSMEG_0250 were purified from *MSMEG mc^2^155* expressing native MSMEG_0736 and MSMEG_0250 with a C-terminal msfGFP-tag. *MSMEG* strains were grown in 7H9 with 0.05% (v/v) Tween_80_. Harvested cells were washed three times with PBS and frozen before lysis with a cryogenic grinder (SPEX SamplePrep). The powder was resuspended in 50 mM Tris-HCl pH 7.5, 150 mM NaCl, protease inhibitor cocktail (Sigma-Aldrich) and 0.6-0.7 units/ml Benzonase endonuclease and the solution was incubated for 30 min. Solutions were centrifuged at 15,000 g for 30 min, followed by centrifugation at 98,000 g - 99,594 g (depending on amount of material) for 1 h to isolate the membranes. Membranes were solubilized for 1 h at 4°C in 50 mM Tris-HCl pH 7.5, 150 mM NaCl and 1% DDM using a 1:10 (w/w) ratio of detergent to membranes. MSMEG_0410 was solubilized using a 1:6.77 (w/w) ratio of detergent to membranes. The solutions were centrifuged for 30 min at 99,526 g - 103,530 g. Solubilized membranes were incubated with GFP-Trap_MA beads (Chromotek) for 1 h at 4°C. The beads were washed three times with 50 mM Tris-HCl pH 7.5, 150 mM NaCl and 0.2% DDM. Proteins were eluted from the beads by the addition of 0.2 M glycine pH 2.5 and the eluate was neutralized with 1 M Tris base pH 10.4. The elution was repeated a second time.

### Mass spectrometry

The two GFP-Trap_MA elutions were pooled and proteins were precipitated with trichloroacetic acid. The pellets were resuspended in 0.1% Rapigest in 50 mM ammonium bicarbonate. Samples were prepared for mass spectrometry analysis as previously described [57]. Samples were denatured and reduced in a buffer containing 2M urea and 2 mM DTT. Free cysteines were alkylated by addition of 2 mM iodoacetamide. The reduced and alkylated samples were then digested with trypsin overnight at 37C. Digested samples were desalted using UltraMicroSpin C18 columns (Nest Group) and then evaporated to dryness. Samples were resuspended in 0.1% formic acid for mass spectrometry analysis.

Samples were analysed on a Thermo Scientific Orbitrap Fusion mass spectrometry system equipped with an Easy nLC 1200 ultra-high pressure liquid chromatography system interfaced via a nanoelectrospray source. Samples were injected onto a C18 reverse phase capillary column (75 um inner diameter x 25 cm length, packed with 1.9 um C18 particles). Peptides were then separated by an organic gradient from 5% to 30% ACN in 0.1% formic acid over 180 minutes at a flow rate of 300 nl/min. The MS continuously collected spectra in a data-dependent fashion over the entire gradient.

Raw mass spectrometry data were analyzed using the MaxQuant software package (version 1.3.0.5) [58]. Data were matched to the *Mycobacterium smegmatis* UniProt reference proteome database. Variable modifications were allowed for methionine oxidation, and protein N-terminus acetylation. A fixed modification was indicated for cysteine carbamidomethylation. Full trypsin specificity was required. The first search was performed with a mass accuracy of +/- 20 parts per million and the main search was performed with a mass accuracy of +/- 6 parts per million. A maximum of 5 modifications were allowed per peptide. A maximum of 2 missed cleavages were allowed. The maximum charge allowed was 7+. Individual peptide mass tolerances were allowed. For MS/MS matching, a mass tolerance of 0.5 Da was allowed and the top 6 peaks per 100 Da were analyzed. MS/MS matching was allowed for higher charge states, water and ammonia loss events. Data were searched against a concatenated database containing all sequences in both forward and reverse directions with reverse hits indicating the false discovery rate of identifications. The data were filtered to obtain a peptide, protein, and site-level false discovery rate of 0.01. The minimum peptide length was 7 amino acids.

Protein identification from a single SDS-PAGE band was performed by the Taplin Mass Spectrometry Facility at Harvard Medical School. The gel band corresponding to the molecular weight of MmpL3 was excised from the gel and subjected to in-gel trypsin digestion. Excised gel bands were cut into approximately 1 mm^3^ pieces. Gel pieces were then subjected to a modified in-gel trypsin digestion procedure [59]. Gel pieces were washed and dehydrated with acetonitrile for 10 min. followed by removal of acetonitrile. Pieces were then completely dried in a speed-vac. Rehydration of the gel pieces was with 50 mM ammonium bicarbonate solution containing 12.5 ng/µl modified sequencing-grade trypsin (Promega, Madison, WI) at 4°C. After 45 min., the excess trypsin solution was removed and replaced with 50 mM ammonium bicarbonate solution to just cover the gel pieces. Samples were then placed in a 37°C room overnight. Peptides were later extracted by removing the ammonium bicarbonate solution, followed by one wash with a solution containing 50% acetonitrile and 1% formic acid. The extracts were then dried in a speed-vac (~1 hr). The samples were then stored at 4°C until analysis.

On the day of analysis the samples were reconstituted in 5 - 10 µl of HPLC solvent A (2.5% acetonitrile, 0.1% formic acid). A nano-scale reverse-phase HPLC capillary column was created by packing 2.6 µm C18 spherical silica beads into a fused silica capillary (100 µm inner diameter x ~30 cm length) with a flame-drawn tip [60]. After equilibrating the column each sample was loaded via a Famos auto sampler (LC Packings, San Francisco CA) onto the column. A gradient was formed and peptides were eluted with increasing concentrations of solvent B (97.5% acetonitrile, 0.1% formic acid).

As peptides eluted they were subjected to electrospray ionization and then entered into an LTQ Orbitrap Velos Pro ion-trap mass spectrometer (Thermo Fisher Scientific, Waltham, MA). Peptides were detected, isolated, and fragmented to produce a tandem mass spectrum of specific fragment ions for each peptide. Peptide sequences (and hence protein identity) were determined by matching protein databases with the acquired fragmentation pattern by the software program, Sequest (Thermo Fisher Scientific, Waltham, MA) [61]. All databases include a reversed version of all the sequences and the data was filtered to between a one and two percent peptide false discovery rate.

### GFPTrap pulldowns of MsTtfA Truncations and MsTtfA-msfGFP with CRISPRi depletion

10ml of LBsmeg culture of MsTfA-msfGFP truncations and MmpL3-mCherry co-expression strains were grown to OD_600_ 0.5 overnight at 37C. 50ml of LBsmeg of TtfA-msfGFP truncations and MmpL3-mCherry co-expression strains with CRISPRi targeting contructs were grown to OD_600_ 0.5. For non-targeting and MSMEG_5308 depletion strains were grown with ATc-50ng/ml for 24 hours and Pks13 depletion strains were grown with ATc for 6 hours. Cultures were cooled on ice and cells were harvested by centrifugation (3700g, 10 min, 4°C). Pellets were washed once with 1ml of PBS. Pellets were resuspended in 500ul PBS with 1x HALT protease (Thermo Scientific) and lysed via bead beating (Biospec, Mini-beadbeater-16) 2 times for 1 min with 5 min on ice between. Beads, unbroken cells, and debris were pelleted at 5000g for 10 min at 4°C. Supernatant was collected and an additional 500ul of PBS containing either 1% DDM or 1% Triton X-100 was added and incubated at 4°C for 1 hour with rocking. Insoluble material was then pelleted at 21130g for 1 hour at 4°C and the supernatant (~1ml) was collected and added to 20ul pre-washed GFPTrap magnetic agarose beads (Bulldog Bio) and incubated for 2 hours at 4°C with rocking. After incubation beads were collected with a magnet and washed 3 times with 1mL PBS and 0.1% DDM or Triton X-100. Elution was done using SDS sample buffer and heating 60°C for 15 min.

## Supporting information

Merged SI tables

## Acknowledgements

This work is supported by AI-U19-111143 (the Tri-I TBRU, part of the TBRU-Network, R01 AI120694, P01 AI063302, P30 CA008748, 5R01AI128214, 1U19AI135990, and P01AI095208.

## Bibliography

1. Banerjee A, Dubnau E, Quemard A, Balasubramanian V, Um KS, et al. (1994) inhA, a gene encoding a target for isoniazid and ethionamide in Mycobacterium tuberculosis. Science 263: 227–230.

2. Goude R, Amin AG, Chatterjee D, Parish T (2009) The arabinosyltransferase EmbC is inhibited by ethambutol in Mycobacterium tuberculosis. Antimicrob Agents Chemother 53: 4138–4146.

3. Telenti A, Philipp WJ, Sreevatsan S, Bernasconi C, Stockbauer KE, et al. (1997) The emb operon, a gene cluster of Mycobacterium tuberculosis involved in resistance to ethambutol. Nat Med 3: 567–570.

4. Bergeret F, Gavalda S, Chalut C, Malaga W, Quemard A, et al. (2012) Biochemical and structural study of the atypical acyltransferase domain from the mycobacterial polyketide synthase Pks13. J Biol Chem 287: 33675–33690.

5. Gavalda S, Bardou F, Laval F, Bon C, Malaga W, et al. (2014) The polyketide synthase Pks13 catalyzes a novel mechanism of lipid transfer in mycobacteria. Chem Biol 21: 1660–1669.

6. Lea-Smith DJ, Pyke JS, Tull D, McConville MJ, Coppel RL, et al. (2007) The reductase that catalyzes mycolic motif synthesis is required for efficient attachment of mycolic acids to arabinogalactan. J Biol Chem 282: 11000–11008.

7. Pacheco SA, Hsu FF, Powers KM, Purdy GE (2013) MmpL11 protein transports mycolic acid-containing lipids to the mycobacterial cell wall and contributes to biofilm formation in Mycobacterium smegmatis. J Biol Chem 288: 24213–24222.

8. Barkan D, Hedhli D, Yan HG, Huygen K, Glickman MS (2012) Mycobacterium tuberculosis lacking all mycolic acid cyclopropanation is viable but highly attenuated and hyperinflammatory in mice. Infect Immun 80: 1958–1968.

9. Barkan D, Liu Z, Sacchettini JC, Glickman MS (2009) Mycolic acid cyclopropanation is essential for viability, drug resistance, and cell wall integrity of Mycobacterium tuberculosis. Chem Biol 16: 499–509.

10. Glickman MS (2003) The mmaA2 gene of Mycobacterium tuberculosis encodes the distal cyclopropane synthase of the alpha-mycolic acid. J Biol Chem 278: 7844–7849.

11. Glickman MS, Cahill SM, Jacobs WR, Jr. (2001) The Mycobacterium tuberculosis cmaA2 gene encodes a mycolic acid trans-cyclopropane synthetase. J Biol Chem 276: 2228–2233.

12. Glickman MS, Cox JS, Jacobs WR, Jr. (2000) A novel mycolic acid cyclopropane synthetase is required for cording, persistence, and virulence of Mycobacterium tuberculosis. Mol Cell 5: 717–727.

13. Huang CC, Smith CV, Glickman MS, Jacobs WR, Jr., Sacchettini JC (2002) Crystal structures of mycolic acid cyclopropane synthases from Mycobacterium tuberculosis. J Biol Chem 277: 11559–11569.

14. Rao V, Gao F, Chen B, Jacobs WR, Jr., Glickman MS (2006) Trans-cyclopropanation of mycolic acids on trehalose dimycolate suppresses Mycobacterium tuberculosis-induced inflammation and virulence. J Clin Invest 116: 1660–1667.

15. Domenech P, Reed MB, Barry CE, 3rd (2005) Contribution of the Mycobacterium tuberculosis MmpL protein family to virulence and drug resistance. Infect Immun 73: 3492–3501.

16. Converse SE, Mougous JD, Leavell MD, Leary JA, Bertozzi CR, et al. (2003) MmpL8 is required for sulfolipid-1 biosynthesis and Mycobacterium tuberculosis virulence. Proc Natl Acad Sci U S A 100: 6121–6126.

17. Bernut A, Viljoen A, Dupont C, Sapriel G, Blaise M, et al. (2016) Insights into the smooth-to-rough transitioning in Mycobacterium bolletii unravels a functional Tyr residue conserved in all mycobacterial MmpL family members. Mol Microbiol 99: 866–883.

18. Szekely R, Cole ST (2016) Mechanistic insight into mycobacterial MmpL protein function. Mol Microbiol 99: 831–834.

19. Sassetti CM, Boyd DH, Rubin EJ (2003) Genes required for mycobacterial growth defined by high density mutagenesis. Mol Microbiol 48: 77–84.

20. Sassetti CM, Rubin EJ (2003) Genetic requirements for mycobacterial survival during infection. Proc Natl Acad Sci U S A 100: 12989–12994.

21. Stec J, Onajole OK, Lun S, Guo H, Merenbloom B, et al. (2016) Indole-2-carboxamide-based MmpL3 Inhibitors Show Exceptional Antitubercular Activity in an Animal Model of Tuberculosis Infection. J Med Chem 59: 6232–6247.

22. Lun S, Guo H, Onajole OK, Pieroni M, Gunosewoyo H, et al. (2013) Indoleamides are active against drug-resistant Mycobacterium tuberculosis. Nat Commun 4: 2907.

23. Rao SP, Lakshminarayana SB, Kondreddi RR, Herve M, Camacho LR, et al. (2013) Indolcarboxamide is a preclinical candidate for treating multidrug-resistant tuberculosis. Sci Transl Med 5: 214ra168.

24. La Rosa V, Poce G, Canseco JO, Buroni S, Pasca MR, et al. (2012) MmpL3 is the cellular target of the antitubercular pyrrole derivative BM212. Antimicrob Agents Chemother 56: 324–331.

25. Xu Z, Meshcheryakov VA, Poce G, Chng SS (2017) MmpL3 is the flippase for mycolic acids in mycobacteria. Proc Natl Acad Sci U S A 114: 7993–7998.

26. Grzegorzewicz AE, Pham H, Gundi VA, Scherman MS, North EJ, et al. (2012) Inhibition of mycolic acid transport across the Mycobacterium tuberculosis plasma membrane. Nat Chem Biol 8: 334–341.

27. Varela C, Rittmann D, Singh A, Krumbach K, Bhatt K, et al. (2012) MmpL genes are associated with mycolic acid metabolism in mycobacteria and corynebacteria. Chem Biol 19: 498–506.

28. Zhang B, Li J, Yang X, Wu L, Zhang J, et al. (2019) Crystal Structures of Membrane Transporter MmpL3, an Anti-TB Drug Target. Cell 176: 636–648 e613.

29. Viljoen A, Dubois V, Girard-Misguich F, Blaise M, Herrmann JL, et al. (2017) The diverse family of MmpL transporters in mycobacteria: from regulation to antimicrobial developments. Mol Microbiol 104: 889–904.

30. Costa TR, Felisberto-Rodrigues C, Meir A, Prevost MS, Redzej A, et al. (2015) Secretion systems in Gram-negative bacteria: structural and mechanistic insights. Nat Rev Microbiol 13: 343–359.

31. Koronakis V, Sharff A, Koronakis E, Luisi B, Hughes C (2000) Crystal structure of the bacterial membrane protein TolC central to multidrug efflux and protein export. Nature 405: 914–919.

32. Du D, Wang Z, James NR, Voss JE, Klimont E, et al. (2014) Structure of the AcrAB-TolC multidrug efflux pump. Nature 509: 512–515.

33. Daury L, Orange F, Taveau JC, Verchere A, Monlezun L, et al. (2016) Tripartite assembly of RND multidrug efflux pumps. Nat Commun 7: 10731.

34. Murakami S, Nakashima R, Yamashita E, Yamaguchi A (2002) Crystal structure of bacterial multidrug efflux transporter AcrB. Nature 419: 587–593.

35. Murakami S, Nakashima R, Yamashita E, Matsumoto T, Yamaguchi A (2006) Crystal structures of a multidrug transporter reveal a functionally rotating mechanism. Nature 443: 173–179.

36. Seeger MA, Schiefner A, Eicher T, Verrey F, Diederichs K, et al. (2006) Structural asymmetry of AcrB trimer suggests a peristaltic pump mechanism. Science 313: 1295–1298.

37. Fu J, Zong G, Zhang P, Gu Y, Cao G (2018) Deletion of the beta-Propeller Protein Gene Rv1057 Reduces ESAT-6 Secretion and Intracellular Growth of Mycobacterium tuberculosis. Curr Microbiol 75: 401–409.

38. Pang X, Cao G, Neuenschwander PF, Haydel SE, Hou G, et al. (2011) The beta-propeller gene Rv1057 of Mycobacterium tuberculosis has a complex promoter directly regulated by both the MprAB and TrcRS two-component systems. Tuberculosis (Edinb) 91 Suppl 1: S142–149.

39. Haydel SE, Clark-Curtiss JE (2006) The Mycobacterium tuberculosis TrcR response regulator represses transcription of the intracellularly expressed Rv1057 gene, encoding a seven-bladed beta-propeller. J Bacteriol 188: 150–159.

40. Manganelli R, Voskuil MI, Schoolnik GK, Smith I (2001) The Mycobacterium tuberculosis ECF sigma factor sigmaE: role in global gene expression and survival in macrophages. Mol Microbiol 41: 423–437.

41. Degiacomi G, Benjak A, Madacki J, Boldrin F, Provvedi R, et al. (2017) Essentiality of mmpL3 and impact of its silencing on Mycobacterium tuberculosis gene expression. Sci Rep 7: 43495.

42. Griffin JE, Gawronski JD, Dejesus MA, Ioerger TR, Akerley BJ, et al. (2011) High-resolution phenotypic profiling defines genes essential for mycobacterial growth and cholesterol catabolism. PLoS Pathog 7: e1002251.

43. Fay A, Glickman MS (2014) An essential nonredundant role for mycobacterial DnaK in native protein folding. PLoS Genet 10: e1004516.

44. Rock JM, Hopkins FF, Chavez A, Diallo M, Chase MR, et al. (2017) Programmable transcriptional repression in mycobacteria using an orthogonal CRISPR interference platform. Nat Microbiol 2: 16274.

45. Feilmeier BJ, Iseminger G, Schroeder D, Webber H, Phillips GJ (2000) Green fluorescent protein functions as a reporter for protein localization in Escherichia coli. J Bacteriol 182: 4068–4076.

46. Yang ZR, Thomson R, McNeil P, Esnouf RM (2005) RONN: the bio-basis function neural network technique applied to the detection of natively disordered regions in proteins. Bioinformatics 21: 3369–3376.

47. Gavalda S, Leger M, van der Rest B, Stella A, Bardou F, et al. (2009) The Pks13/FadD32 crosstalk for the biosynthesis of mycolic acids in Mycobacterium tuberculosis. J Biol Chem 284: 19255–19264.

48. Leger M, Gavalda S, Guillet V, van der Rest B, Slama N, et al. (2009) The dual function of the Mycobacterium tuberculosis FadD32 required for mycolic acid biosynthesis. Chem Biol 16: 510–519.

49. Carel C, Nukdee K, Cantaloube S, Bonne M, Diagne CT, et al. (2014) Mycobacterium tuberculosis proteins involved in mycolic acid synthesis and transport localize dynamically to the old growing pole and septum. PLoS One 9: e97148.

50. Tahlan K, Wilson R, Kastrinsky DB, Arora K, Nair V, et al. (2012) SQ109 targets MmpL3, a membrane transporter of trehalose monomycolate involved in mycolic acid donation to the cell wall core of Mycobacterium tuberculosis. Antimicrob Agents Chemother 56: 1797–1809.

51. Domenech P, Reed MB, Dowd CS, Manca C, Kaplan G, et al. (2004) The role of MmpL8 in sulfatide biogenesis and virulence of Mycobacterium tuberculosis. J Biol Chem 279: 21257–21265.

52. Jain M, Cox JS (2005) Interaction between polyketide synthase and transporter suggests coupled synthesis and export of virulence lipid in M. tuberculosis. PLoS Pathog 1: e2.

53. Rath P, Huang C, Wang T, Wang T, Li H, et al. (2013) Genetic regulation of vesiculogenesis and immunomodulation in Mycobacterium tuberculosis. Proc Natl Acad Sci U S A 110: E4790–4797.

54. Fulop V, Jones DT (1999) Beta propellers: structural rigidity and functional diversity. Curr Opin Struct Biol 9: 715–721.

55. Chaudhuri I, Soding J, Lupas AN (2008) Evolution of the beta-propeller fold. Proteins 71: 795–803.

56. Chen CK, Chan NL, Wang AH (2011) The many blades of the beta-propeller proteins: conserved but versatile. Trends Biochem Sci 36: 553–561.

57. Jager S, Cimermancic P, Gulbahce N, Johnson JR, McGovern KE, et al. (2011) Global landscape of HIV-human protein complexes. Nature 481: 365–370.

58. Cox J, Mann M (2008) MaxQuant enables high peptide identification rates, individualized p.p.b.-range mass accuracies and proteome-wide protein quantification. Nat Biotechnol 26: 1367–1372.

59. Shevchenko A, Wilm M, Vorm O, Mann M (1996) Mass spectrometric sequencing of proteins silver-stained polyacrylamide gels. Anal Chem 68: 850–858.

60. Peng J, Gygi SP (2001) Proteomics: the move to mixtures. J Mass Spectrom 36: 1083–1091.

61. Eng JK, McCormack AL, Yates JR (1994) An approach to correlate tandem mass spectral data of peptides with amino acid sequences in a protein database. J Am Soc Mass Spectrom 5: 976–989.

