## Supplementary figures and images for "Two accessory proteins govern MmpL3 mycolic acid transport in mycobacteria"

### SI figures

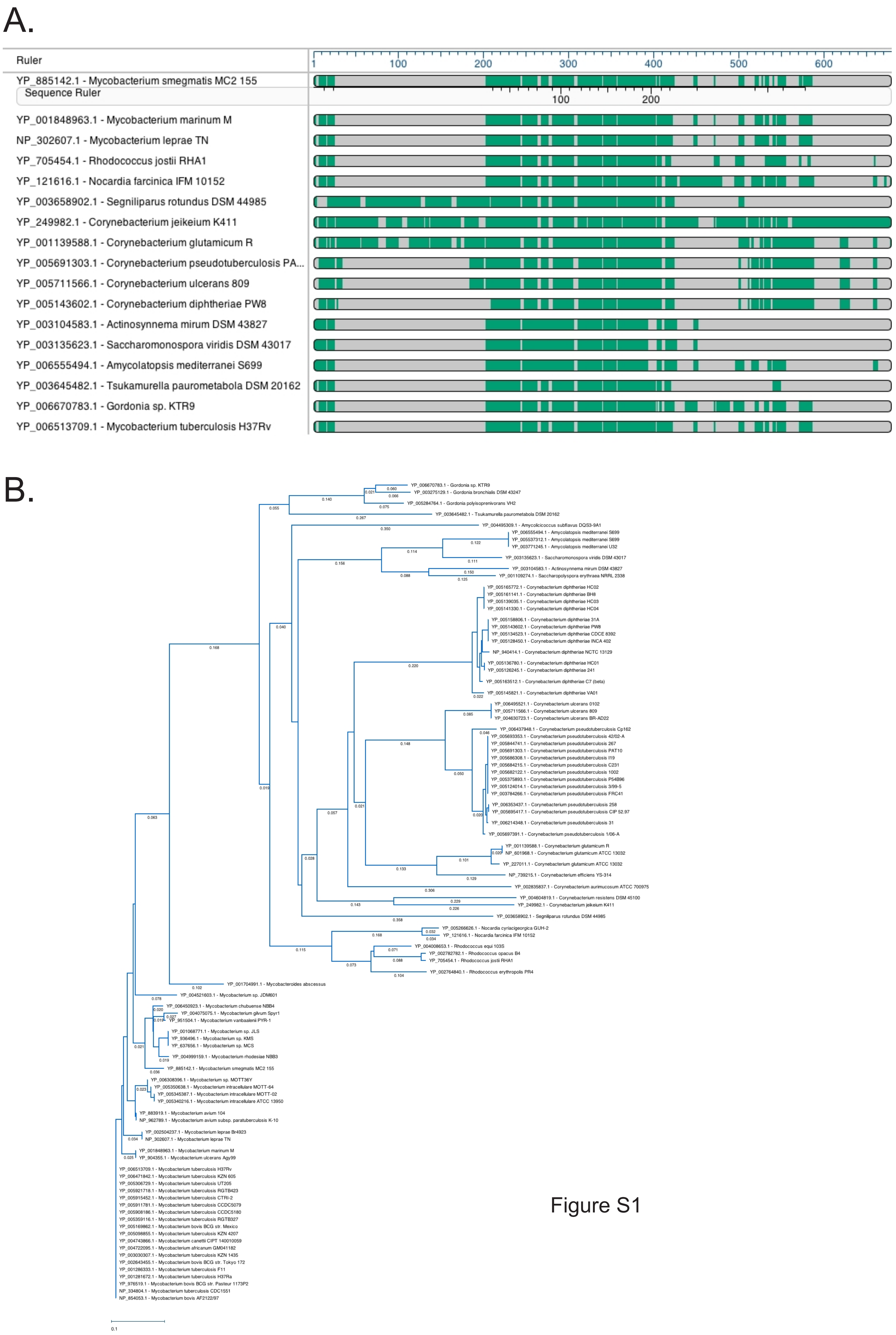

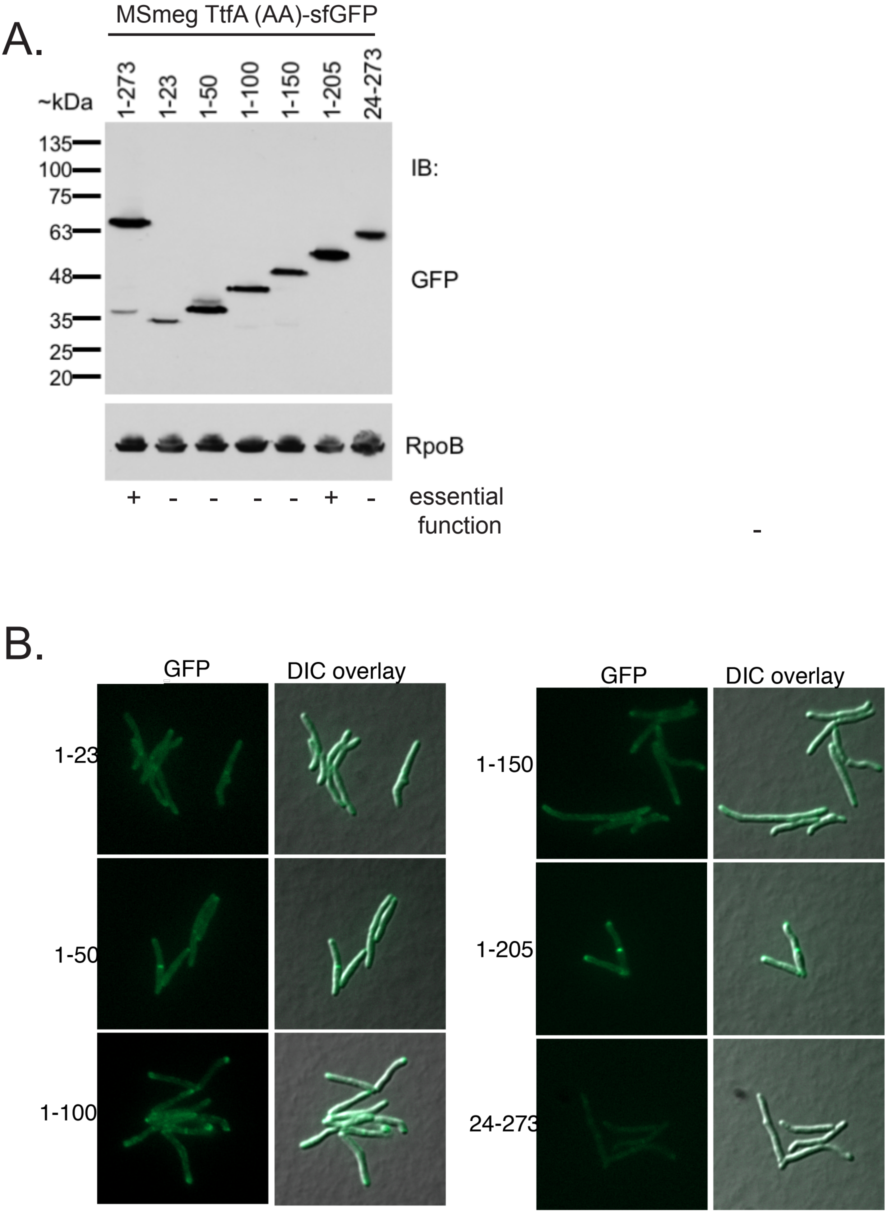

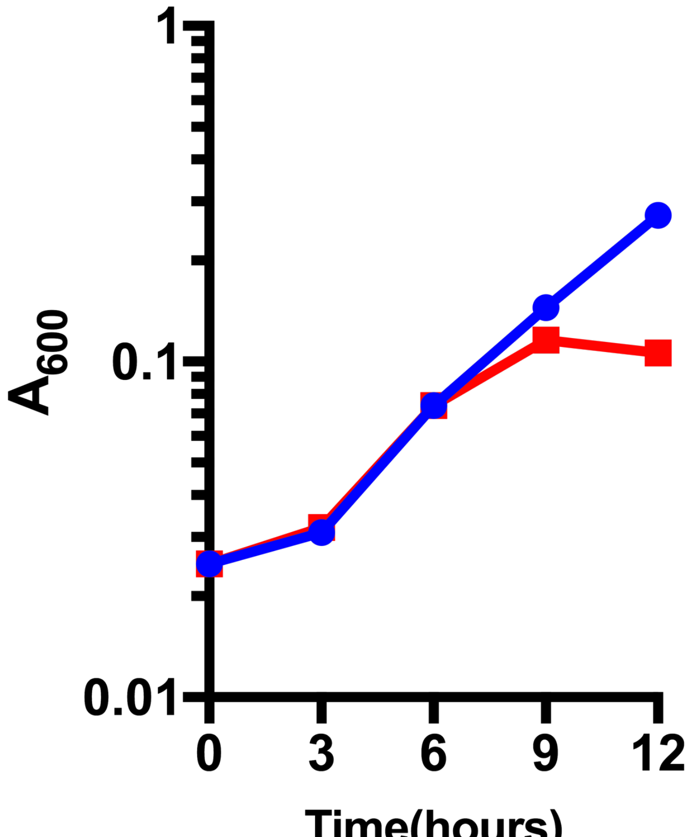


Figure S3

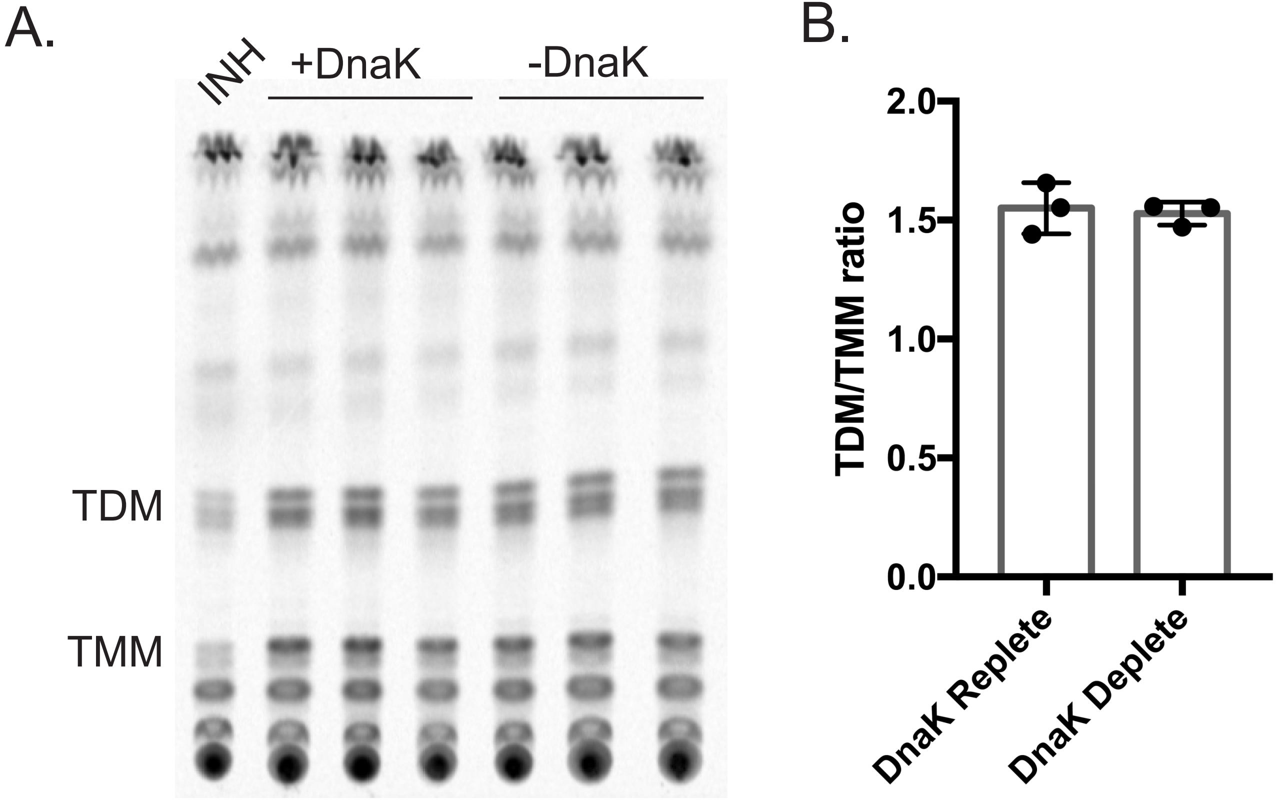

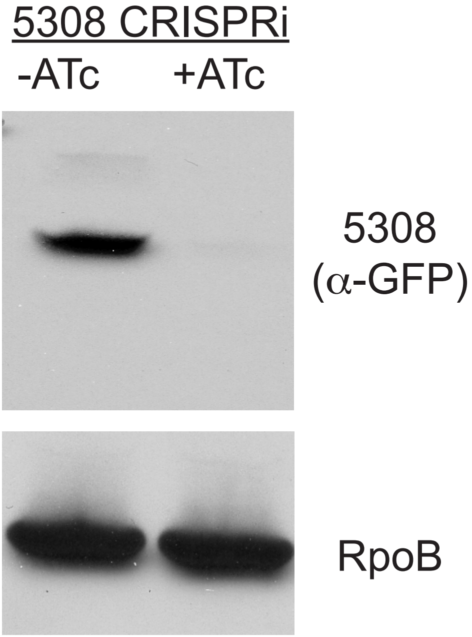


Figure S5
